# In Silico Analysis and Characterization of Differentially Expressed Genes to Distinguish Glioma Stem Cells from Normal Neural Stem Cells

**DOI:** 10.1101/2021.04.05.438487

**Authors:** Urja Parekh, Mohit Mazumder, Harpreet Kaur, Elia Brodsky

**Affiliations:** Deparment of Biological Science, Sunandan Divatia School of Science, Narsee Monjee Institute of Management Studies, India; Pine Biotech, USA

**Keywords:** RNA-Seq Analysis, DGE, Machine Learning, Glioblastoma multiforme, Cancer stem cells

## Abstract

Glioblastoma multiforme (GBM) is a heterogeneous, invasive primary brain tumor that develops chemoresistance post therapy. Theories regarding the aetiology of GBM focus on transformation of normal neural stem cells (NSCs) to a cancerous phenotype or tumorigenesis driven via glioma stem cells (GSCs). Comparative RNA-Seq analysis of GSCs and NSCs can provide a better understanding of the origin of GBM. Thus, in the current study, we performed various bioinformatics analyses on transcriptional profiles of a total 40 RNA-seq samples including 20 NSC and 20 GSC, that were obtained from the NCBI-SRA (SRP200400). First, differential gene expression (DGE) analysis using DESeq2 revealed 348 significantly differentially expressed genes between GSCs and NSCs (padj. value <0.05, log2fold change ≥ 3.0 (for GSCs) and ≤ −3.0 (for NSCs)) with 192 upregulated and 156 downregulated genes in GSCs in comparison to NSCs. Subsequently, exploratory data analysis using principal component analysis (PCA) based on key significant genes depicted the clear separation between both the groups. Further, Hierarchical clustering confirmed the distinct clusters of GSC and NSC samples. Eventually, the biological enrichment analysis of the significant genes showed their enrichment in tumorigenesis pathways such as Wnt-signalling, VEGF-signalling and TGF-β-signalling pathways. Conclusively, our study depicted significant differences in the gene expression patterns between NSCs and GSCs. Besides, we also identified novel genes and genes previously unassociated with gliomagenesis that may prove to be valuable in establishing diagnostic, prognostic biomarkers and therapeutic targets for GBM.

## Introduction

Glioblastoma multiforme (GBM), a grade IV glioma that accounts for over 60% of all primary brain tumors, is associated with very poor prognosis and an overall survival period of a mere 15 months post-surgery, radiotherapy and temozolomide (TMZ) chemotherapy (Stupp et al., 2009). It is an aggressive and recurring cancer with a median relapse rate of 7 months (Weller et al., 2009). Despite several advances in therapies over the past few years, GBM remains one of the most devastating and difficult brain cancers to treat owing to its interpatient and intratumoral heterogeneity, subsequent resistance to chemotherapy, lack of significant therapeutic biomarkers and inaccessibility of the tumors, for therapeutic intervention, based on their locations in the brain (Holland, 2000; Weller et al., 2009).

Glioblastoma is termed multiforme because its phenotypic and genetic heterogeneity imparts complexity to this particular cancer. GBM tumors have been classified into mesenchymal, classical, proneural/IDH mutant and proneural/RTK mutant variants depending on their molecular signatures (Chen et al., 2017). 90% of GBMs occur de novo in elderly patients, while secondary GBMs arise from low grade astrocytomas in younger patients. Known genetic hallmarks of primary GBM include gene mutation and amplification of epidermal growth factor receptor (EGFR), overexpression of mouse double minute 2 (MDM2), deletion of p16 and loss of heterozygosity (LOH) of chromosome 10q holding phosphatase and tensin homolog (PTEN) and TERT promoter mutation. In contrast, secondary GBM have different genetic signatures such as over expression of platelet-derived growth factor A, and platelet-derived growth factor receptor alpha (PDGFA/PDGFRa), retinoblastoma (RB), LOH of 19q and mutations of IDH1/2, TP53 and ATRX (Hanif et al., 2017). Further, single-cell RNA-Seq studies have revealed that different cell types with varying genetic patterns may be present within a tumor, which may be helpful in determining the prognosis of the tumor. Cancer stem cells account for one such small population of cells and are predicted to drive tumorigenesis and impart chemoresistance in GBM (Couturier et al., 2020).

Despite several efforts made by researchers to understand the cytogenetic aspects of these tumors, the etiology of it still remains largely unknown. Two hypotheses have been postulated that suggest its origin. The cancer stem cell theory purports that GBM originates from cancer stem cells (CSCs) that are responsible for self-renewal, development, propagation and recurrence of the tumor. The other theory states that GBM arises via transformation of normal neural stem cells after accumulation of several mutations in common neural stem cell marker genes (Couturier et al., 2020; Yao et al., 2018). This concept is rather complex since GSCs may share certain genetic similarities with NSCs, however, the molecular differences may underpin the malignant growth potential of the tumor. Thus, it is important to understand the differences in gene expression between normal neural stem cells (NSCs) and Glioblastoma Stem Cells (GSCs) to be able to better determine the cellular origin of the tumor.

In this study, we performed RNA-Seq analysis using NSC and GSC samples to understand the differences in their gene expression profiles, understand the origin of GBM, and identify potential biomarkers that may allow for selective targeting of CSCs while sparing normal NSCs via precision medicine.

## Methods

### Acquisition of RNA-Seq Data

A thorough search of the Gene Expression Omnibus (GEO) Database was performed to obtain the ideal dataset for the study. The GSE132172 (associated Sequence Retrieval Archive (SRA) Study: SRP200400) dataset was selected (Zhao et al., 2019). This dataset consisted of RNA-Seq data retrieved from CB660 normal neural stem cell lines and GliNS2 glioblastoma stem cell lines. Of the 188 samples present on the associated SRA Run Selector, 20 samples of NSCs (SRR9200813 to SRR9200832) and 20 samples of GSCs (SRR9200895 to SRR9200914) were selected and downloaded as an SRA Run Table.

### RNA-Seq Data Analysis

PCR clean and Trimmomatic algorithms were used to remove adapter sequence and low quality sequence data, mapping on genome and junctions was done via TopHat2, isoform construction via cufflinks, Gene transfer format (GTF) processing via cuffmerge, mapping on transcripts via Bowtie2-t and the final gene expression values were obtained using RSEM algorithm as RsemExpTable in Fragments Per Kilobase of transcript per Million mapped reads (FPKM) units (figure 1). The gene expression data was log transformed and quantile normalized for further analysis.

**Figure 1.**
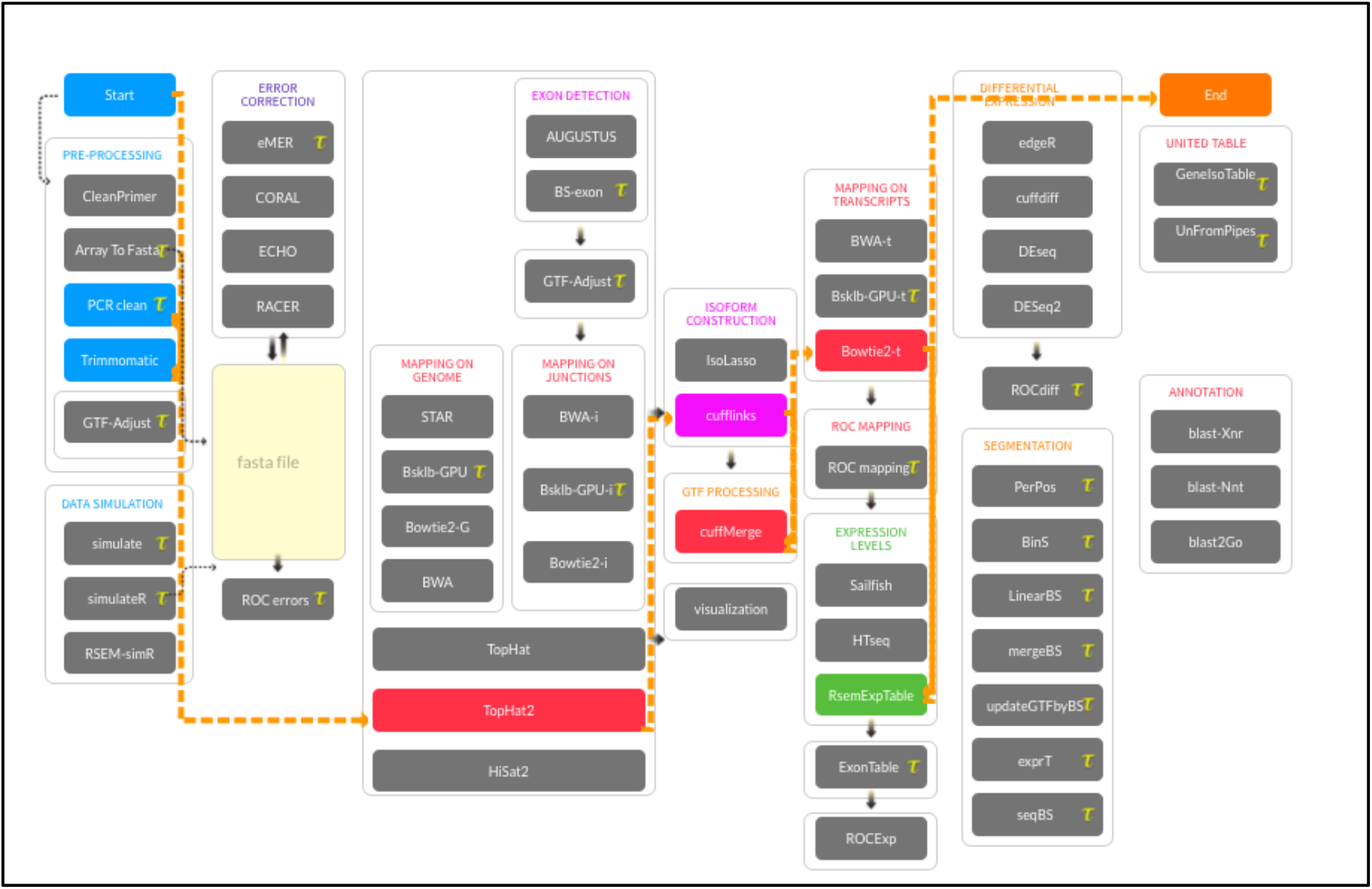
Screenshot of RNA-Seq Pipeline (Tuxedo Protocol).

### Downstream Data Analysis

The 40 samples were clustered as per their gene expression profiles using Principal Component Analysis and Hierarchical Clustering (distance: Euclidean, linkage: ward.D2). Differential gene expression was performed using the DESeq2 pipeline to derive significantly expressed genes in GSC and NSC samples (figure 2). The differential gene expression data was then filtered and extracted if the threshold = TRUE, p-adjusted value was <0.05 and log_2_fold change value for GSCs was ≥ 3.0 (for GSCs) and ≤ −3.0 (for NSCs). Top 25 most highly expressed genes from each type of stem cell sample was further filtered and a heat map and dendrogram was generated to depict comparison of gene expression profiles in each type of sample.

**Figure 2.**
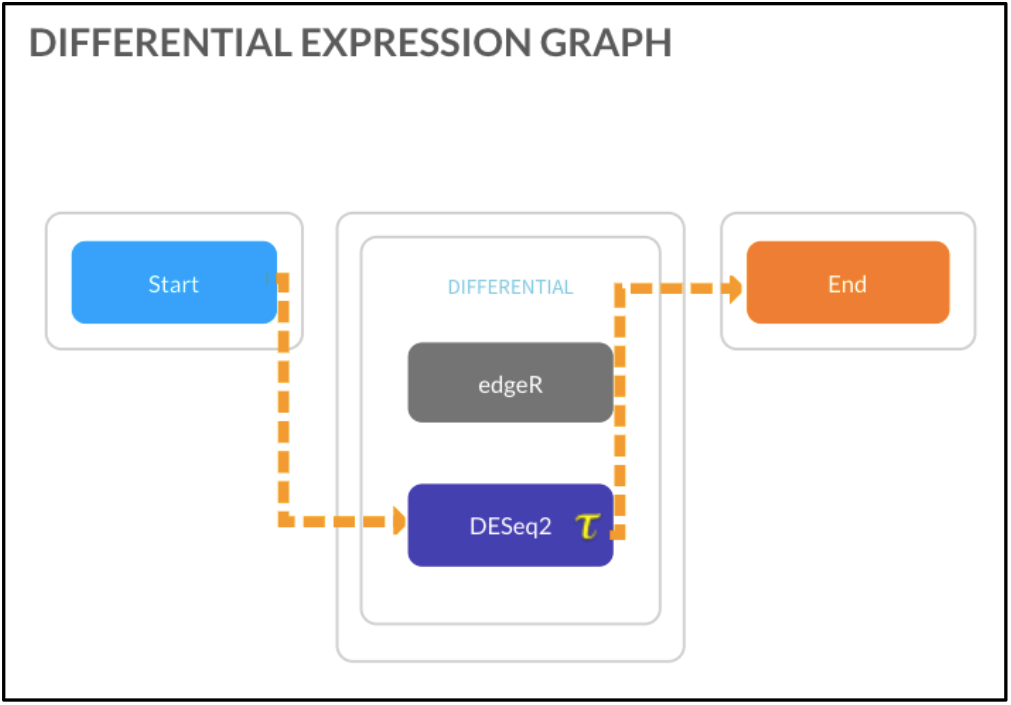
Screenshot of differential gene expression analysis using DeSeq2 Pipeline.

### Gene Enrichment Analysis

To understand the biological implication of the significant genes obtained from differential gene expression analysis, the gene lists were uploaded on the Database for Annotation, Visualization and Integrated Discovery (DAVID) v6.8 tool. Kyoto Encyclopedia of Genes and Genomes (KEGG) was also used for pathway analysis. Literature review for the top 50 most significant differentially expressed genes (which include top 25 upregulated and top 25 down regulated genes in GSCs vs. NSCs) was performed via Google Scholar, National Center for Biotechnology Information (NCBI) PubMed and GeneCards®, to understand their functions in a normal physiological condition and in gliomagenesis.

## Results

Gene expression data in FPKM units for a total of 27,385 genes was obtained in the RSemExp table after the RNA-Seq Pipeline (Tuxedo Protocol) was run. Post quantile normalization and log-scale transformation, PCA plot was generated which revealed separate clustering of GSC and NSC samples with a principal component 1 (PC1) of 10.31% and PC2 of 8.9% (supplementary figure 1a). A single outlier (SRR9200898_PE), which was a glioma stem cell sample, was revealed. Thus, another PCA plot was generated without the outlier. A PC1 of 88.03% and a PC2 of 2.13% was obtained with this plot (supplementary figure 1b). To further confirm these findings, hierarchical clustering was performed that revealed NSC samples (SRR9200813 to SRR9200832) and GSC samples (SRR9200895 to SRR9200897 and SRR9200899 to SRR9200914) clustered separately. The outlier GSC sample (SRR9200898_PE) lay between the two clusters (supplementary figure 2).

Further analysis done via differential gene expression revealed 12,437 (45.42% of the total genes) differentially expressed genes between NSC and GSC samples. This data was filtered and a total of 951 significantly expressed genes (7.64% of all differentially expressed genes) with a threshold = TRUE and p-adjusted value ≤ 0.05 was obtained. A volcano plot was generated to depict these significantly differentially expressed genes between NSCs and GSCs (figure 3). Further filtering was performed to narrow down the most highly differentially expressed genes between NSCs and GSCs. A threshold of ≥ 3.0 (for GSCs) and ≤ −3.0 (for NSCs) was set for the log_2_fold change value. Thus, we were left with 348 significantly differentially expressed genes. Among them, 192 genes were found to be upregulated and 156 to be downregulated in GSCs in comparison to NSCs. Further, we selected top 50 significantly differentially expressed genes including top 25 upregulated and top 25 downregulated genes in GSCs vs. NSCs (Table 1)

**Table 1:**
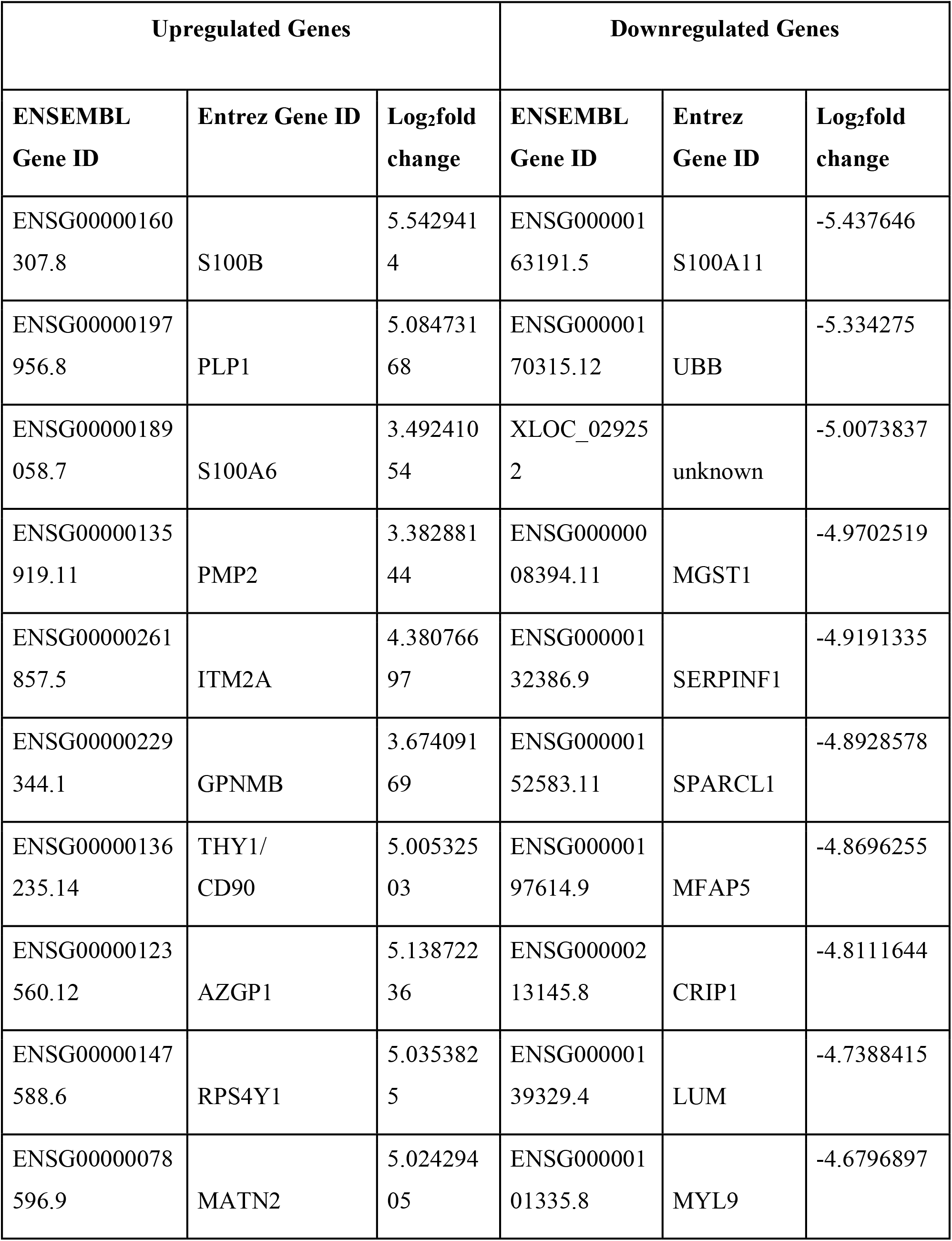
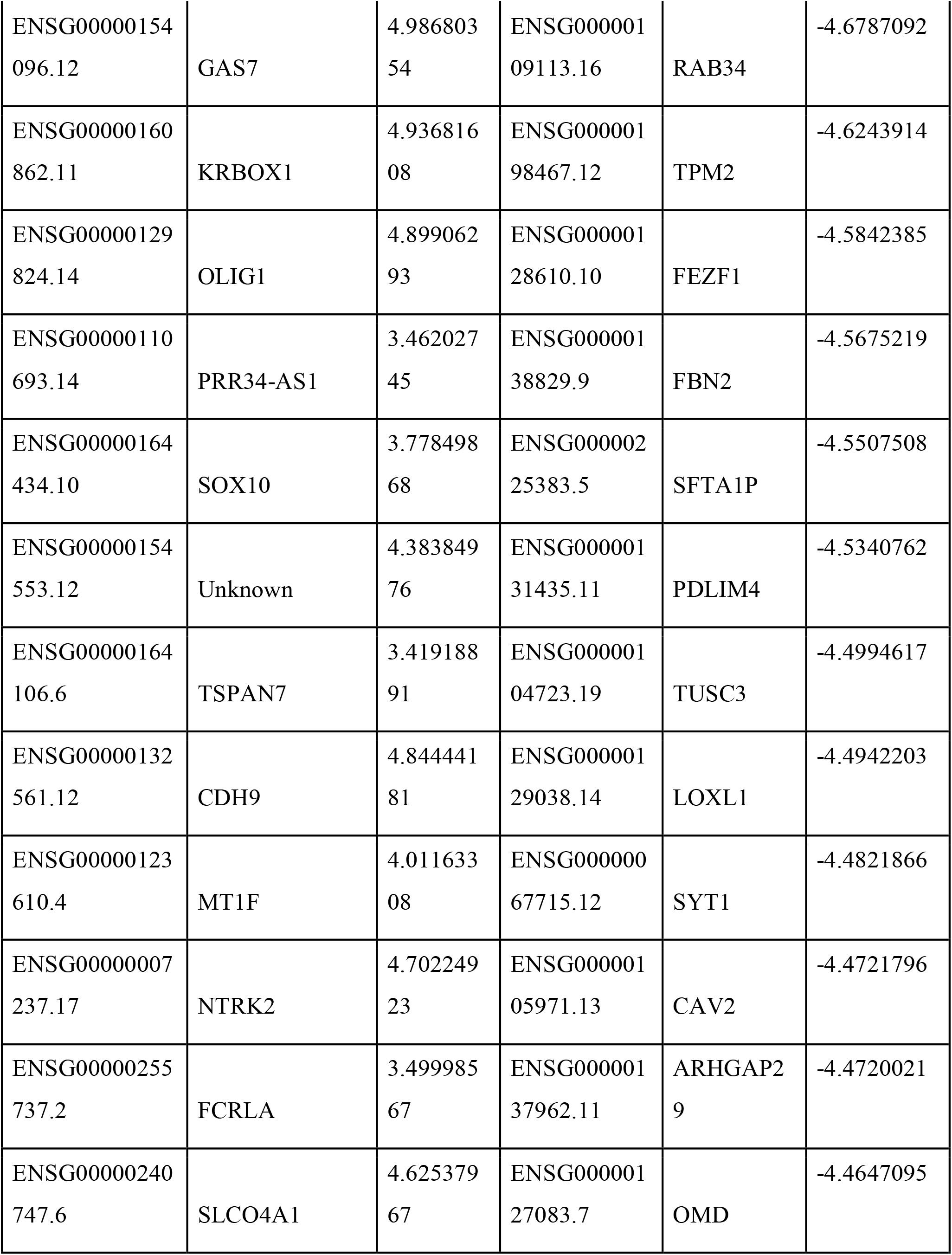
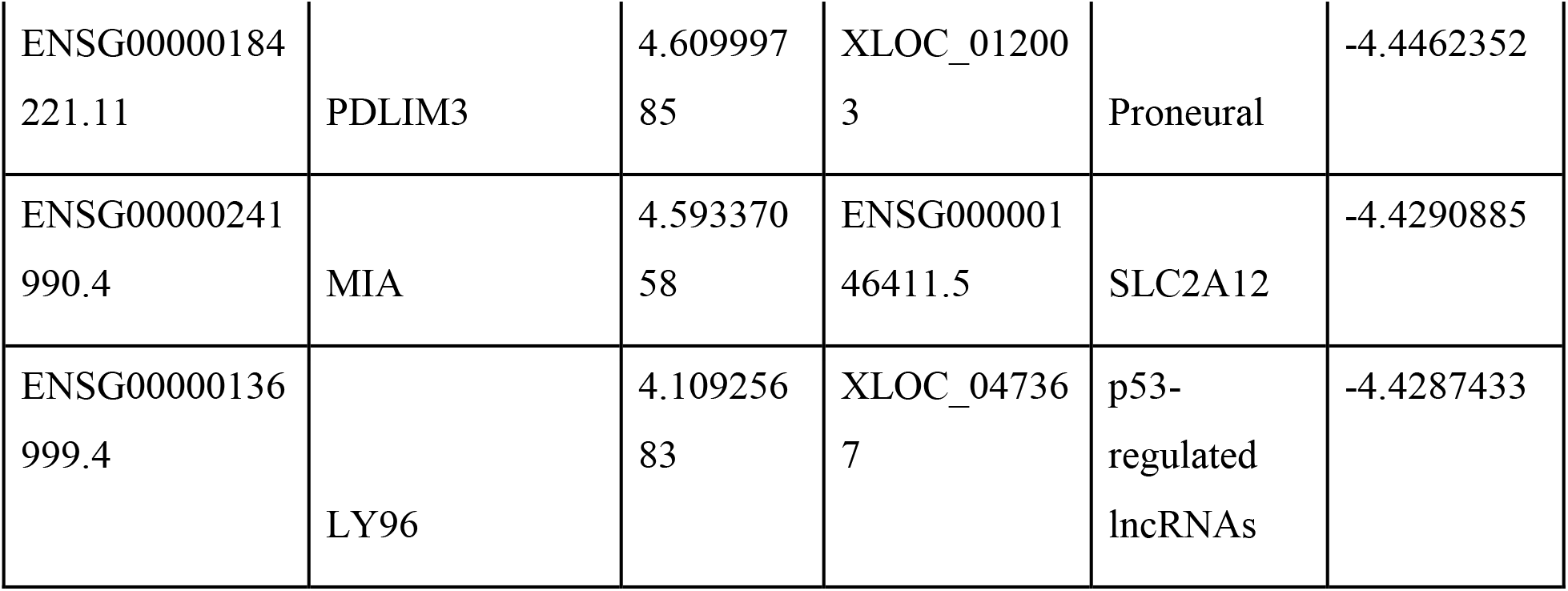
Top 50 differentially expressed genes in GSCs vs NSCs

**Figure 3.**
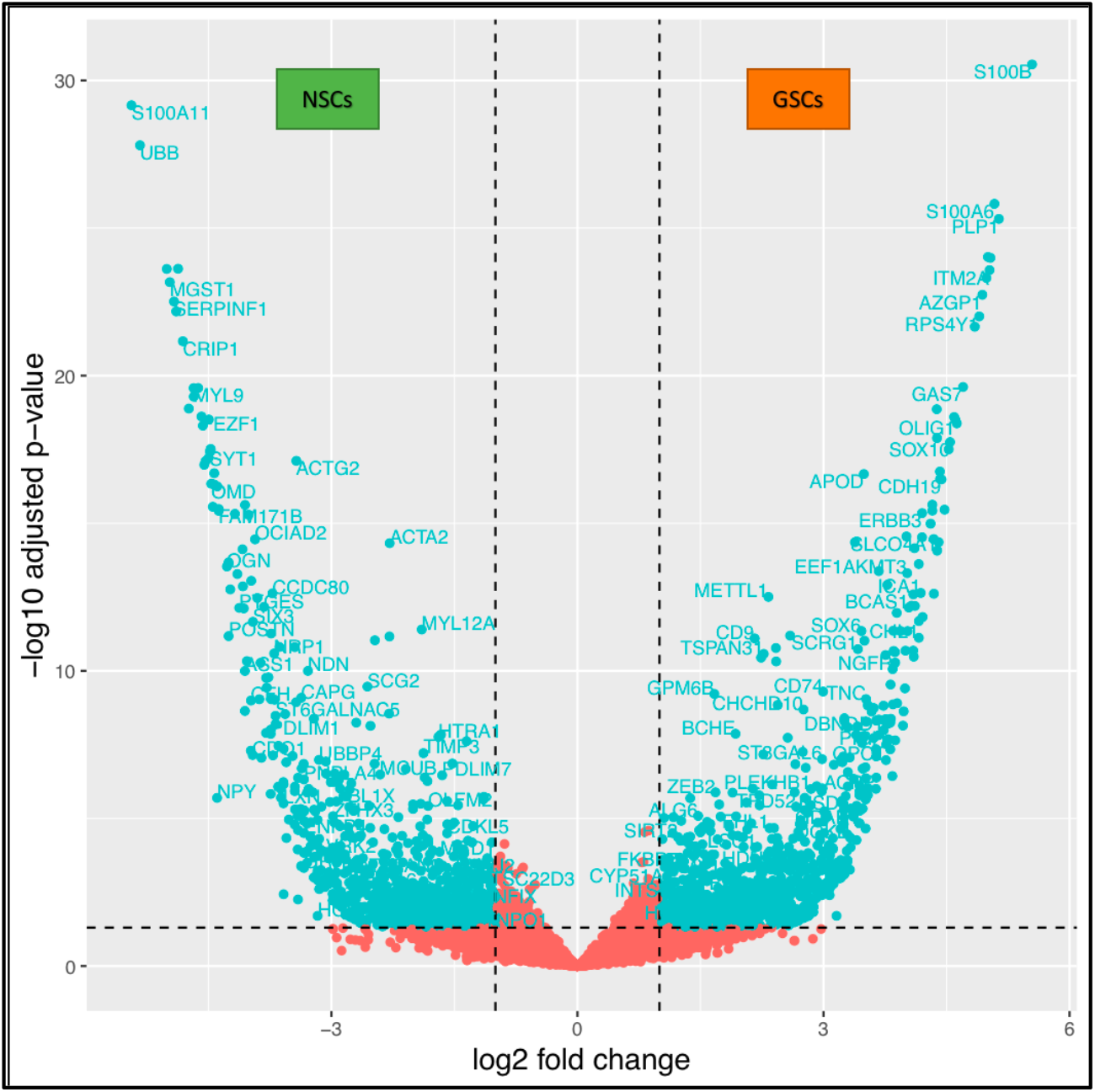
Volcano plot of differentially expressed genes between GSCs and NSCs.

Next, to determine the distinguishable potential of these top 50 significantly differentially expressed genes, we generated the PCA plots with only these significant genes with the outlier (Figure 4a) and without the outlier (Figure 4b). Subsequently, in the heatmap (Figure 5), we visualized the differential expression pattern of these genes among two types of samples, i.e. GSCs vs NSCs.

**Figure 4.**
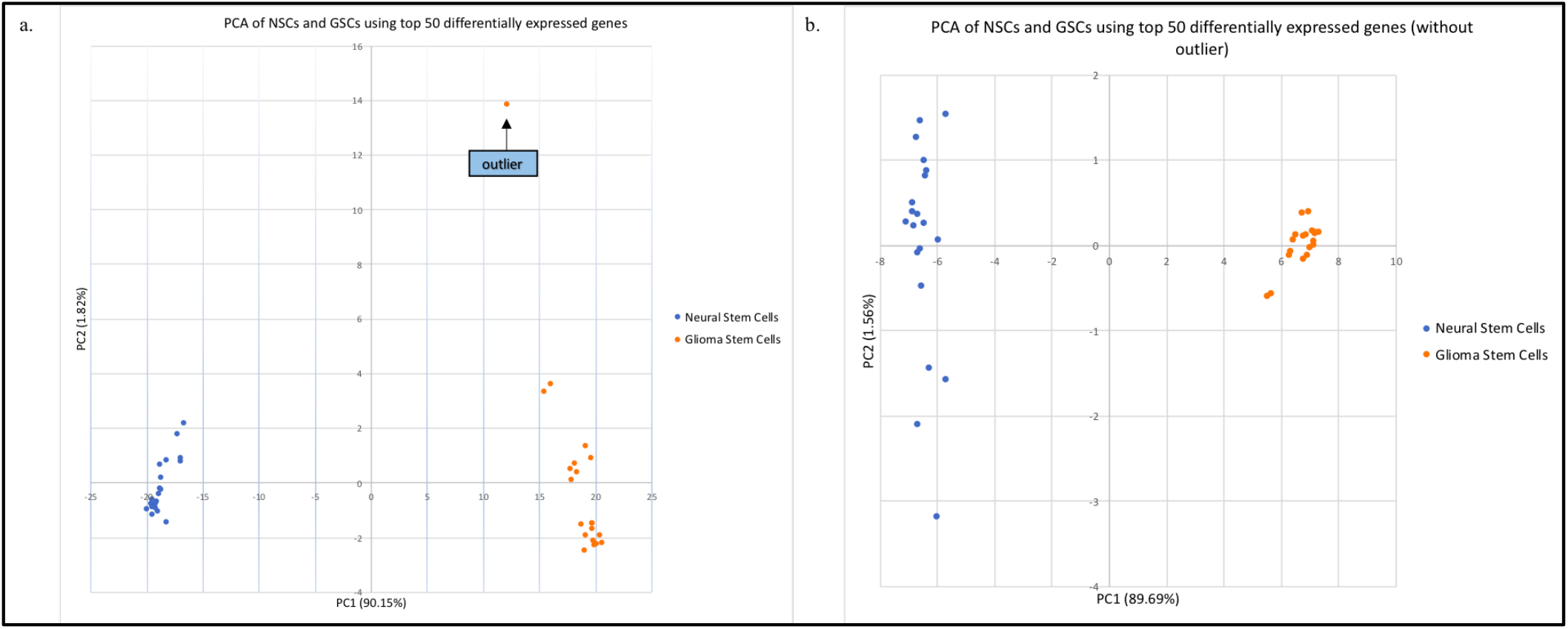
Exploratory data analysis using PCA based on the top 50 significantly differentially expressed genes between NSCs and GSCs. (a). PCA plot for all GSC and NSC samples, (b) PCA plot for GSC and NSC samples after removal of outlier.

**Figure 5.**
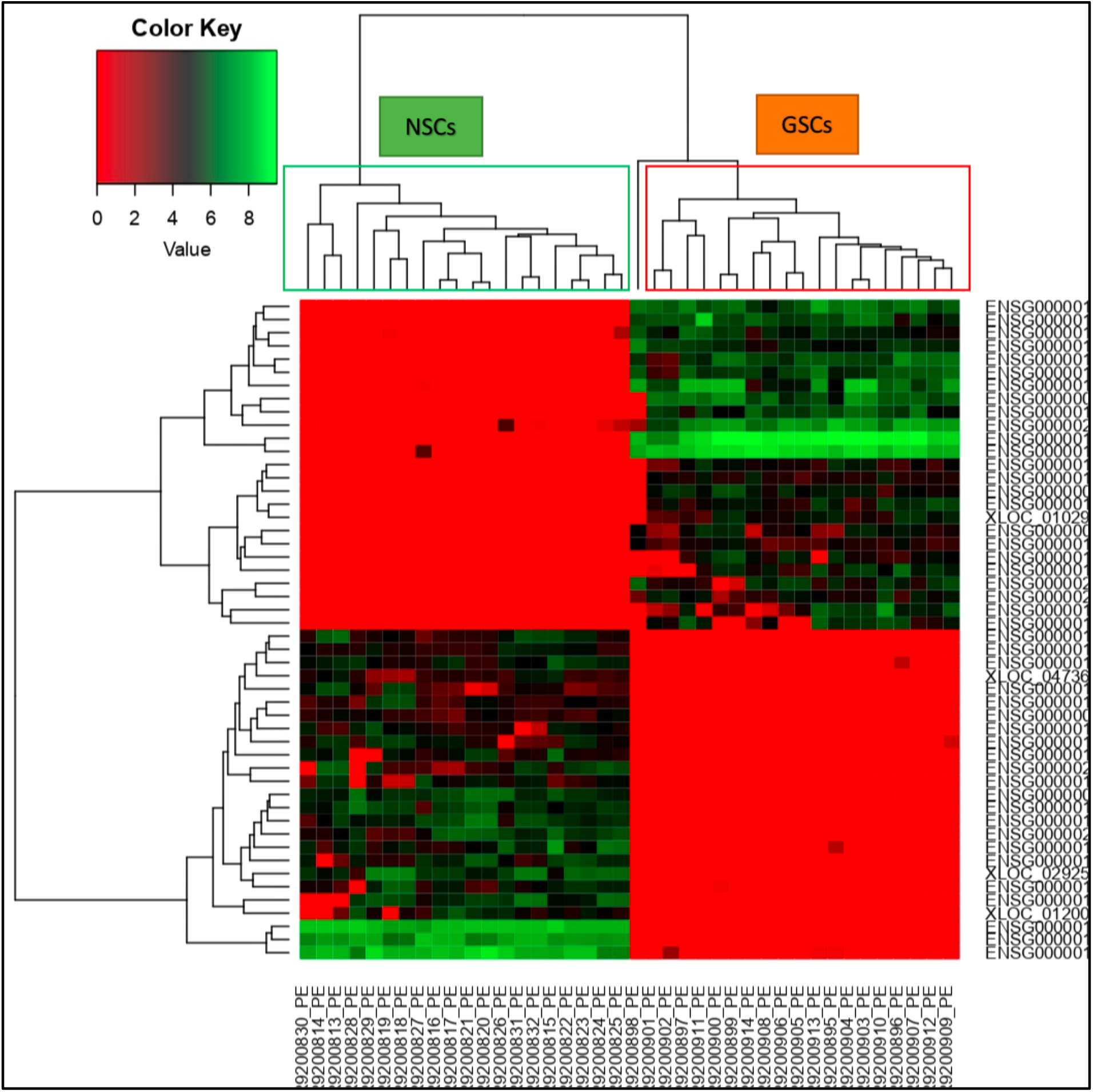
Heat map representing the expression pattern of the top 50 differentially expressed genes expressed genes between NSCs and GSCs.

Finally, we re-performed H-Clustering with the top 50 selected genes (figure 6). The GSC and NSC samples clustered separately, with the outlier sample branching in between the two clusters.

**Figure 6.**
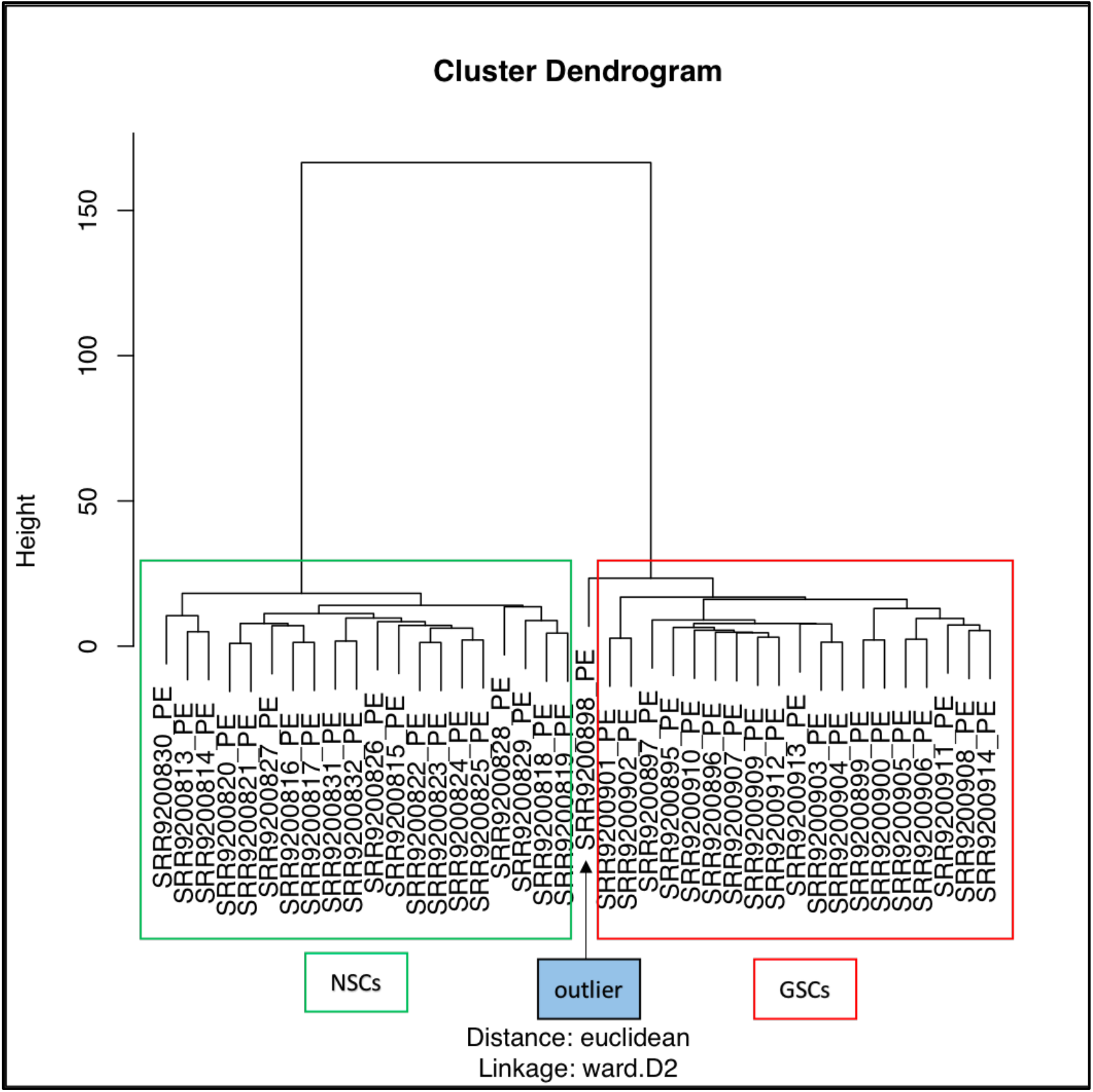
Dendrogram generated by Hierarchical clustering based on top 50 significantly differentially expressed genes depicting the distinct clusters of GSCs and NSCs samples.

### Gene Enrichment analysis

The gene list for NSC containing 156 genes and for GSC containing 192 genes was uploaded on DAVID and functional annotation and clustering was performed.

Top hits with the NSC gene list indicated involvement in diseases such as Alzheimer’s Disease, respiratory function tests, Tobacco Use Disorder, body weight, bone mineral density, macular degeneration, alcoholism, etc. Top hits for keyword annotations included extracellular matrix, secreted, glycoprotein, signal, calcium, disulphide bond, disease mutation, membrane and transmembrane. Top hits for sequence features included signal peptide, glycosylation (N-linked), Leucine Rich Repeats (LRR 6), EGF-like 6, LRR 5, LRR 1, LRR 2, LRR 7, LRR 4, disulphide bond, topological domain (cytoplasmic). Functional clustering (medium stringency) showed 27 clusters with 137 DAVID IDs with top three clusters involved in extracellular matrix, glycoprotein and LRRs and having enrichment scores of 5.99, 4.95 and 2.05, respectively.

Similarly, the gene list for GSC containing 192 genes was also uploaded on DAVID and top hits for diseases included Schizophrenia, Attention deficit hyperactivity disorder (ADHD), tobacco use disorder, depression, body height, body weight, autism, myocardial Infarction. Top hits for keyword annotations included disulphide bond, glycoprotein, signal, MHC-II, cell adhesion, cell membrane, alternative splicing, polymorphism, transmembrane. Top hits for sequence features included glycosylation (N-linked), signal peptide, disulphide bond, topological domain (extracellular and cytoplasmic), transmembrane, splice variant, sequence variant. Functional clustering (medium stringency) showed 27 clusters with 163 DAVID IDs with top three clusters involved in glycoprotein disulphide bonds, melanocyte differentiation metallothionein domain and having enrichment scores of 6.17, 2.35 and 2.15, respectively.

KEGG Pathway analysis depicted genes such as Ankyrin, α2β1, Lumican, Wnt, α5β1, SDC-4 and Caveolin from the list that were involved in various tumorigenesis pathways. Other genes such as PVRL3, PVRL2, NCAM, L1CAM, IGSF4, CDH3 and NEO1, enriched in the neural system, were also depicted. ECM Receptor Interaction genes such as Fibronectin, Collagen, Laminin, Tenascin, αV, β8, Thrombospondin (THBS) and Osteopontin (OPN), were observed in the pathway analysis. Genes such as FAK, PAK, RasGAP, Ephrin A, Ephrin B, Robo1, Robo3, ERK, Cofilin, GSK3β, Plexin B involved in axon guidance pathways, and genes such as PLCβ, PLCγ, RTK, PMCA involved in calcium signaling pathways were also enriched in the gene lists (figure 7).

**Figure 7:**
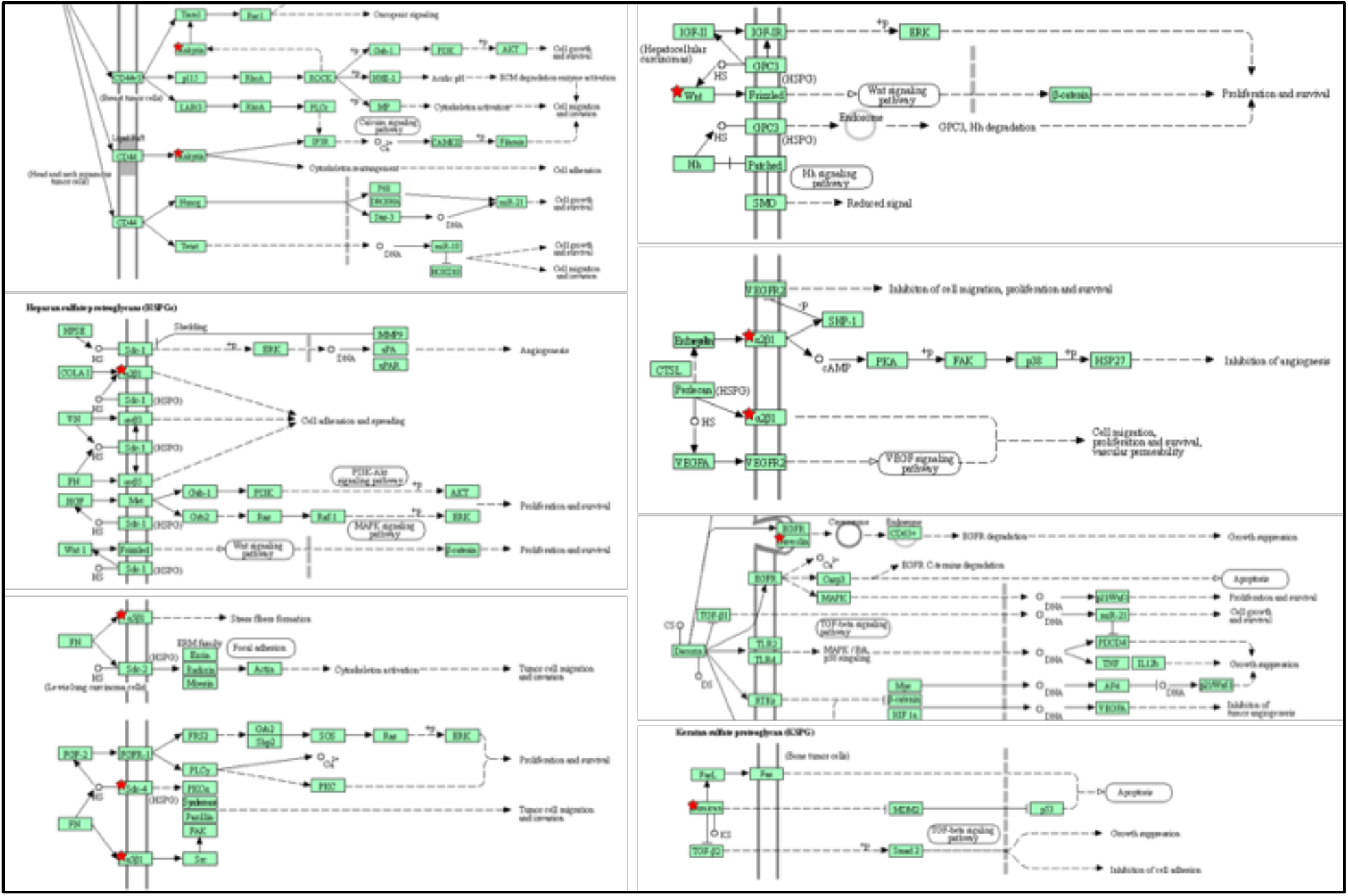
Pathway analysis of significant genes in GSC and NSC samples.

The detailed information regarding the roles and functions of the top 25 highly upregulated and 25 highly downregulated genes in GSCs and NSCs, within normal pathophysiology and gliomagenesis is tabulated in Tables (see Supplementary tables S1 and S2). Upregulated genes such as S100B, S100A6, GPNMB, CD90, SOX10 to name a few have established association with gliomagenesis, whereas genes such as ITM2A, KRBOX1, PRR34-AS1, SLCO4A1 and novel gene ENSG00000154553 have no known association with the disease (Avril et al., 2017, p. 90; Donato et al., 2017; Ferletta et al., 2007, p. 10; Frauchiger et al., 2019; Liguori et al., 2020). Similarly downregulated genes with established association are S100A11, UBB, MGST1, SERPINF1 and SPARCL1, while no previous association has been established for XLOC_02 9252 (novel gene), FBN2, SFTA1P, OMD, XLOC_01 2003 (novel gene), XLOC_04 7367 (novel gene) (Armento et al., 2017; Gagliardi et al., 2020; Kedves et al., 2017; Tu et al., 2019, p. 11; Xu et al., 2017).

## Discussion

The origin of GBM from cancer stem cells or via transformation of normal neural stem cells, still remains largely unknown. Previous studies addressing this question have shown contrasting results, thus making it difficult to establish any one of the theories as the answer (Couturier et al., 2020; Mukherjee, 2020; Yao et al., 2018; Zhao et al., 2019). In this study we have made an attempt to reveal the origin of GBM via RNA-Seq analysis.

To identify differences in gene expression between normal NSCs and GSCs, RNA-Seq analysis was performed on RNA-seq samples obtained from both cell populations. Analysis revealed that both cell populations clustered separately on PCA, depicting the vast genetic differences between the two populations of stem cells. Further, removal of the outlier while performing PCA increased the variance contributed by principal component (PC1) significantly representing substantial variability across samples. The outlier GSC sample, that clustered closer to normal NSCs (also shown in the cluster dendrogram), is of utmost importance as it indicates that the outlier could have genetic similarities with both NSCs and GSCs. This hints towards the theory that GBM may develop via accumulation of mutations in the normal NSCs thus transforming to GSCs. Further studies on the same would be able to divulge the series of events that take place during this transformation.

Based on differential gene expression analysis using DESeq2, we observed 192 genes that were significantly (adjusted P-value <0.05, log2 fold change ≥ 3.0 (for GSCs) and ≤ −3.0 (for NSCs)) upregulated and 156 genes that were downregulated in GSCs in comparison to NSCs. Furthermore, to get a manageable gene set, we selected only the top 50 genes, i.e. top 25 upregulated and top 25 downregulated genes with highest fold change values in GSC vs. NSCs samples. Next, the PCA plot and dendrogram from Hierarchical clustering indicated their significance in determining the differences between GSCs and NSCs. A heat map generated from these genes showed that most genes upregulated in GSCs were downregulated in NSCs and vice versa, highlighting differential expression in both populations.

Gene ontology analysis performed on DAVID using significant gene sets showed no obvious involvement of genes, specifically in gliomagenesis, via functional annotation and clustering. However, hits were obtained for genes associated with cell-adhesion, migration and invasion, indicating that their possible dysregulation may have led to tumorigenesis. Further, pathway analysis demonstrated a better understanding of the involvement of the gene sets in pathways representing the hallmarks of cancer such as cell growth, survival, proliferation, adhesion, migration, invasion, growth suppression, apoptosis, angiogenesis and vascular permeability. For instance, we observed the activation of the TGF-β pathway. Previous studies have shown that glioma stem cells release TGF-β and activate this notorious signaling pathway, to induce epithelial-mesenchymal transition (EMT) and increase the invasiveness of the glioma (Ye et al., 2012). Activation of the Wnt-signaling pathway was also seen, however, not many studies have defined its role in GBM pathogenesis, and thus, it would be interesting to understand its function in the etiology of this cancer (Guan et al., 2020). Another important pathway identified in our study was the VEGF signaling pathway, a known pathway activated in GSCs, inducing cancer-stem cell proliferation and therefore, a potential therapeutic target for GBM (Xu et al., 2013). Other pathways associated with normal functioning of neurons were also observed that included the calcium signaling, axon guidance and presynaptic-postsynaptic pathways. Previous studies have their involvement in the initiation, development and prognosis of GBM (Li et al., 2019; Yang et al., 2019).

This study was able to identify several target genes whose role in gliomagenesis remains largely unclear. Studying their functionality in the future may reveal key biomarkers or drug targets that could be exploited for GBM.

## Conclusion

Differences in gene expression between glioma stem cells and normal neural stem cells were identified using RNA-Seq analysis. Discovery of novel genes and genes with no known association in gliomagenesis were important outcomes of our study. Studying the outlier (SRR9200898_PE) in further detail would most likely reveal important clues to the etiology of this fatal cancer.

## Limitations of the study

We selected a small sample size of NSCs and GSCs that were derived from a heterogeneous population with unknown demographic features of the patients, and lacked data regarding their treatment status. This prevents the generalizability of our results to a larger population. As part of a future study, it would be interesting to understand the changes in gene expression in both NSCs and GSCs post therapy with current gold standards of treatment for GBM versus new modalities such as nanotherapy and precision drugs like bevacizumab (Gilbert et al., 2014; Grauer et al., 2019; Stupp et al., 2009b).

## Acknowledgements

I would like to thank the entire team at Pine Biotech for their invaluable guidance, training and resources provided for this project. The project was completed during the course of the OmicsLogic Fellowship Program. Pipelines for the study were prepared using the T-BioInfo server.

## Authors’s Contribution

Urja Parekh proposed the research idea, conducted the analysis and wrote the article. Mentorship and guidance was provided by Dr. Mohit Mazumdar and Elia Brodsky through the course of the project. Dr. Harpreet Kaur reviewed and edited the article.

