## Supplementary Data for "In Silico Analysis and Characterization of Differentially Expressed Genes to Distinguish Glioma Stem Cells from Normal Neural Stem Cells"

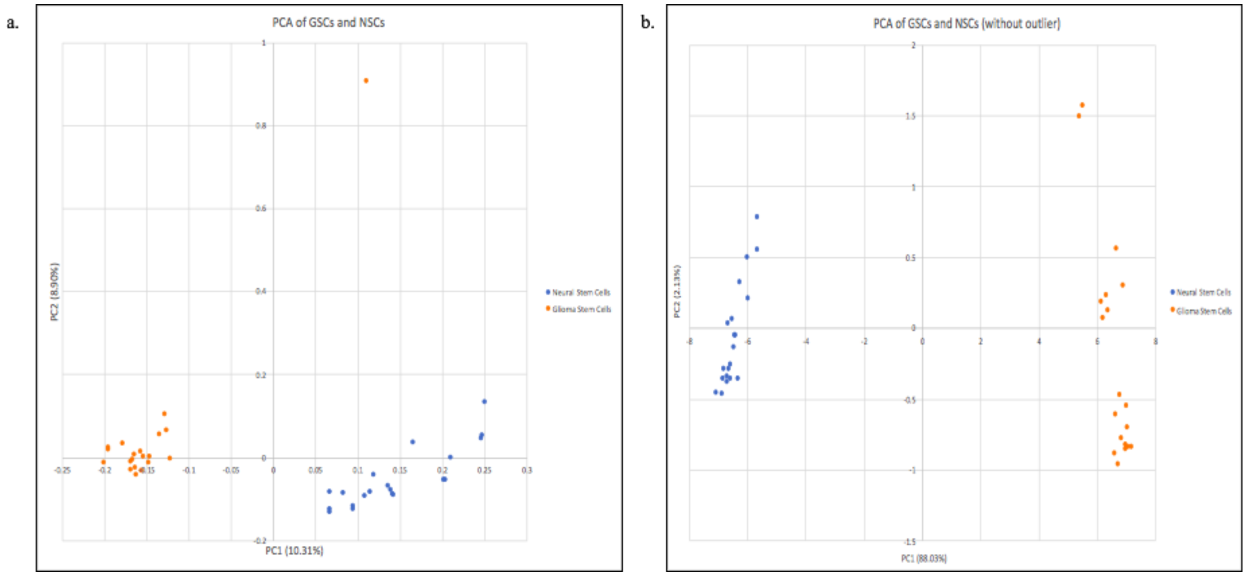


**Supplementary Figure 1.** Exploratory data analysis of NSCs and GSCs samples using PCA based on all the genes: (**a)** PCA plot for all samples, (**b)** PCA plot of all samples after removal of outlier sample (SRR9200898_PE).

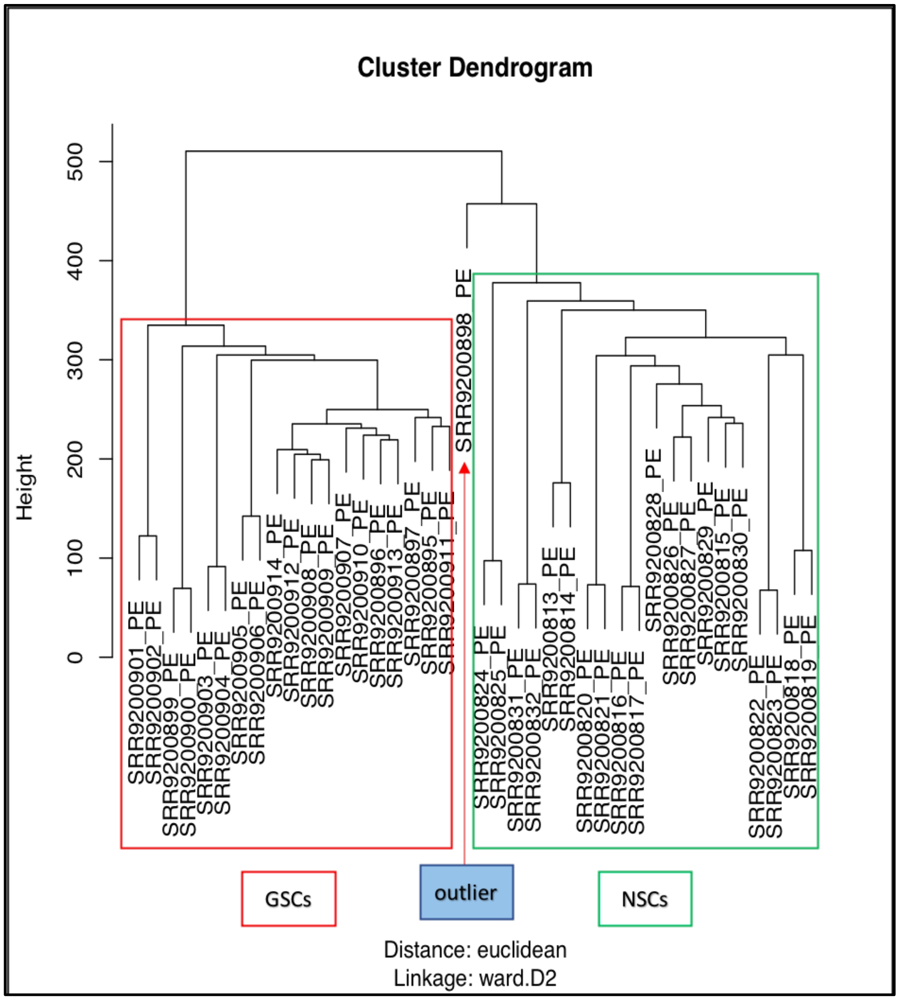


**Supplementary Figure 2.** Dendrogram from H-Clustering analysis depicting the clusters of NSC and GSC samples based on all genes.

**Supplementary Table S1:** List of top 25 upregulated genes with their roles in normal cells and Glioma.

| ENTREZ ID | Full name | Function in normal pathology | Function in glioma | References |
|---|---|---|---|---|
| S100B | S100 Calcium Binding Protein B | This gene encodes a protein with functions in neurite extension, proliferation of melanoma cells, stimulation of Ca2+ fluxes, inhibition of PKC-mediated phosphorylation, astrocytosis and axonal proliferation, and inhibition of microtubule assembly. | Shorter survival period is associated with high levels of serum S100B in glioma patients. It promotes glioma growth by Tumor-associated macrophages (TAMs) chemoattraction through upregulation of CCL2 and thus S100B inhibitors are potential drugs for glioma therapy. | (Frauchiger et al., 2019; Gao et al., 2018; Holla et al., 2016; Sorci et al., 2013) |
| PLP1 | Proteolipid Protein 1 | This gene encodes a transmembrane proteolipid protein present in myelin which play a role in maintaining the compaction and stabilization of myelin sheaths, as well as in | PLP1 association with glioma invasion. Needs further investigation to determine a significant functional role. | (Luo et al., 2019; Miyamoto et al., 2012; Ocklenburg et al., 2019) |

| | | | | |
|---|---|---|---|---|
| | | development and survival of oligodendrocytes. | | |
| S100A6 | S100 Calcium Binding Protein A6 | Involved in the regulation of a number of cellular processes such as cell cycle progression and differentiation. | S100A6 are expressed in a small subset of cancer stem cells and that its overexpression is positively correlated to tumor grade. | (Donato et al., 2017, p. 6; Harris et al., 2008) |
| PMP2 | Peripheral Myelin Protein 2 | The protein encoded by this gene is a component of the myelin sheaths of the peripheral nervous system. Mutation in this gene is shown to be the cause of dominant demyelinating CMT neuropathy. | PMP2 is associated with the stemness of GBM and a discriminatory biomarker between gliomas and gliosarcomas. However, individual gene studies needed. | (Hong et al., 2016, p. 2; Vital et al., 2010) |
| ITM2A | Integral Membrane Protein 2A | In vivo studies have suggested the protein's role in osteo- and chondrogenic differentiation. | Little known. In other cancers it prevents proliferation and invasion. More studies are required. | (Deleersnijder et al., 1996; Zhang et al., 2021, p. 2) |

| | | | | |
|---|---|---|---|---|
| GPNMB | Glycoprotein nonmetastatic melanoma protein B | GPNMB may be responsible for prolonging cell survival and tumor growth while increasing metastatic potential of malignant tumors. | Overexpression of GPNMB is associated with a poor prognosis in glioblastoma patients. GPNMB promotes glioma growth via Na+/K+-ATPase α subunits which can be exploited as a novel therapeutic target for the treatment of GBM. | (Liguori et al., 2020; Ono et al., 2016) |
| THY1/ CD90 | Thy-1 Cell Surface Antigen | The encoded protein is involved in cell adhesion and cell communication in cells of the immune and nervous systems. This gene may function as a tumor suppressor in nasopharyngeal carcinoma. | CD90 is an established stem cell marker and expressed in GBM-associated stromal cells (GASCs) and mesenchymal stem cell-like pericytes, indicative of the heterogenous nature of GBM. It is therefore a good therapeutic target. | (Avril et al., 2017, p. 90; Sauzay et al., 2019) |

| | | | | |
|---|---|---|---|---|
| AZGP1 | Alpha-2-Glycoprotein 1, Zinc-Binding | Stimulates lipid degradation in adipocytes and causes the extensive fat losses associated with some advanced cancers. | It is one of the overexpressed genes in the GBM Extracellular Vesicles protein signature. More studies are needed. | (Huang et al., 2013; Osti et al., 2019) |
| RPS4Y1 | Ribosomal Protein S4 Y-Linked 1 | It is a Y chromosome gene having an important role in prostate cancer. | Associated with GBM but requires more studies | (Khosravi et al., 2014; Wipfler et al., 2018) |
| MATN2 | Matrilin 2 | This gene encodes a extracellular matrix (ECM) protein that balances the communication between ECM and epithelial cells. | Associated with GBM, however, more studies are needed. | (Varga et al., 2010; Zhang et al., 2014) |

| | | | | |
|---|---|---|---|---|
| GAS7 | Growth Arrest Specific 7 | This protein is expressed in terminally differentiated brain cells such as in mature cerebellar Purkinje neurons. It plays an important role in neuronal development. | GAS7 has been detected at low to moderate levels and is not deemed prognostic for glioma. More studies are required. | (Ju et al., 1998, p. 7; Tatenhorst et al., 2005; You and Lin-Chao, 2010, p. 7) |
| KRBOX1 | KRAB Box Domain Containing 1 | KRBOX1 are DNA-binding repressors of transcription. The targets of this protein remain largely unknown. | Has not been associated yet, thus, requires further studies. | (Park et al., 2017) |
| OLIG1 | Oligodendrocyte Transcription Factor 1 | OLIG1 (Oligodendrocyte Transcription Factor 1) is a protein-coding gene associated with Oligodendroglioma and Grade Iii Astrocytoma. It is involved in pathways such as Neural Crest Differentiation and Neural Stem Cells and Lineage-specific Markers. | Lower expression of Olig1 observed in glioblastoma. Olig1 transcription factor is required for maturation of oligodendrocyte progenitors. | (Bouvier et al., 2003, p. 1; Dai et al., 2015; Wu et al., 2012) |

| | | | | |
|---|---|---|---|---|
| PRR34-AS1 | PRR34 Antisense RNA 1 | PRR34-AS1 (PRR34 Antisense RNA 1) is an RNA Gene, and is affiliated with the lncRNA class. It has been identified to play a role in medulloblastoma. | No associated found yet, thus, need more studies. | (Kesherwani et al., 2020) |
| SOX10 | SRY-Box Transcription Factor 10 | It encodes a protein that acts as a nucleocytoplasmic shuttle protein and is important for neural crest and peripheral nervous system development. Among its related pathways are Neural Crest Differentiation and Neural Stem Cells and Lineage-specific Markers. | Sox10 has been identified as a master-regulator of the RTKI subtype in GBM. Low-grade gliomas have a higher expression of Sox10 compared to high-grade glioma. | (Ferletta et al., 2007, p. 1; Pozniak et al., 2010; Rehberg et al., 2002; Wu et al., 2020) |
| ENSG00000154553.12 (unknown) | N/A | N/A | N/A | N/A |
| TSPAN7 | Tetraspanin 7 | This glycoprotein and may have a role in the control of neurite outgrowth. Among its related pathways are dysregulation of transcription in cancer and | TSPAN7 has been identified as a good prognostic biomarker, associated with a longer disease-free | (Bassani et al., 2012; Wuttig et al., 2012) |

| | | | | |
|---|---|---|---|---|
| | | chemical transmission across synapses. | survival and tumor-specific survival. | |
| CDH9 | Cadherin 9 | This gene encodes a type II classical cadherin from the cadherin superfamily, integral membrane proteins that mediate calcium-dependent cell-cell adhesion. It has been associated to play a role in the development of autism spectrum disease in cerebellar pathways. | CDH9 has a role in the transition of CSCs to Endothelial Cells (ECs). More studies are required. | (Turaga and Lathia, 2016; Wang et al., 2019, p. 9) |
| MT1F | Metallothionein 1F | Metallothioneins have a high content of cysteine residues that bind various heavy metals; these proteins are transcriptionally regulated by both heavy metals and glucocorticoids. It may be an essential tumor-growth suppressor in hepatocellular carcinoma and colon cancer. | MT1F has shown to play a role in imparting chemoresistance in glioblastoma cells. It may also be a significant prognostic biomarker to differentiate short- and long-term patient survival. | (Lu et al., 2003; Mehrian-Shai et al., 2015; Yan et al., 2012) |

| | | | | |
|---|---|---|---|---|
| NTRK2 | Neurotrophic Receptor Tyrosine Kinase 2 | This gene encodes a member of the neurotrophic tyrosine receptor kinase (NTRK) family that plays a role in a signalling pathway leading to cell survival and differentiation. | NTRK2 is known to rearrange or fuse with other genes such as BCR and is associated with driving tumorigenesis and increasing aggressiveness of glioblastoma. | (Jones et al., 2019; Pattwell et al., 2020a, 2020b; Torre et al., 2020) |
| FCRLA | Fc Receptor Like A | This protein may also be involved in the development of lymphomas. It has several known spliced transcript isoforms. | No known studies performed yet and thus, further investigation needed. | (Capone et al., 2016; Santiago et al., 2011) |
| SLCO4A1 | Solute Carrier Organic Anion Transporter Family Member 4A1 | SLCO4A1 is involved in several pathways such as transport of vitamins, nucleosides, and related molecules and transport of glucose and other sugars, bile salts and organic acids, metal ions and amine compounds. Its role has been established in several cancers including colorectal cancer. | Has still not been associated with GBM and requires further studies. One study found it to be expressed in differentially methylated region GBM cells. | (Ban et al., 2017; Sun et al., 2018) |

| | | | | |
|---|---|---|---|---|
| PDLIM3 | PDZ And LIM Domain 3 | The protein encoded by this gene contains a PDZ domain and a LIM domain, indicating that it may be involved in cytoskeletal assembly. It is known to be involved in cardiac and muscular disease pathogenesis. | Associated but needs further studies. | (Kim et al., 2013; Maurin et al., 2009) |
| MIA | MIA SH3 Domain Containing | Diseases associated with MIA include Melanoma. Among its related pathways are Neural Crest Differentiation. It is responsible for growth inhibition in melanoma, neuroectodermal tumors, including gliomas. | MIA mRNA expression has been observed in melanoma and glioma tumor cells but not in non-glioma CNS tumors making it a potential diagnostic biomarker. More studies are needed. | (Hau et al., 2002; Poser et al., 2004) |
| LY96 | Lymphocyte Antigen 96 | This gene encodes a protein which associates with toll-like receptor 4 on the cell surface and confers responsiveness to lipopolysaccyaride (LPS), thus providing a link between the receptor and LPS signaling. | Associated with poor survival in GBM, however, needs further studies. | (Dou et al., 2013; Moreno et al., 2021; Rajaraman et al., 2009) |

**Supplementary Table S2**: List of top 25 upregulated genes with their roles in normal cells and gliomas.

| ENTREZ ID | Full name | Function | Function in glioma | References |
|---|---|---|---|---|
| S100A11 | S100 Calcium Binding Protein A11 | This protein plays a role in the functions of motility, invasion, tubulin polymerization. Mutations and chromosomal rearrangements of this gene may play a role in tumor metastasis. | S100A11 is a prognostic marker for GBM with high expression indicating poor outcome. It's overexpression promotes cell proliferation, epithelial-mesenchymal transition (EMT), migration and invasion of glioma stem cells (GSCs), whereas its knockdown has shown to inhibit these roles. | (Mori et al., 2004, p. 1; Tu et al., 2019, p. 11) |
| UBB | Ubiquitin B | This gene encodes ubiquitin, that is involved in the maintenance of chromatin structure, regulation of gene expression, and stress response. Its repression has been linked with ovarian, uterine and endometrial cancers. | Associated with the regulation of several pathways of GBM, however their specific function remains unknown. More studies are required. | (Kedves et al., 2017; Scholz et al., 2020) |
| XLOC_029252 | Novel gene | N/A | N/A | N/A |

| | | | | |
|---|---|---|---|---|
| MGST1 | Microsomal Glutathione S-Transferase 1 | This protein protects the membranes of the endoplasmic reticulum and mitochondria from oxidative stress. This gene is also essential for embryonic development and hematopoiesis in vertebrates. | Downregulation of MGST1 weakens cell adhesion (may lead to metastasis). However, no specific association with GBM has been found, and thus, more studies are required. | (Armento et al., 2017; Bräutigam et al., 2018) |
| SERPINF1 | Serpin Family F Member 1 | It is a strong inhibitor of angiogenesis. It is a neurotrophic factor involved in neuronal differentiation in retinoblastoma cells. | Loss of expression of this gene is involved in glioma proliferation. | (Guan et al., 2004; Xu et al., 2017) |
| SPARCL1 | SPARC Like 1 | It is a tumor suppressor gene in several types of tumor while it may also be an oncogene in other types. | This gene is a potential therapeutic biomarker for GBM as it may be involved in driving the growth and infiltration of GSCs while also promoting angiogenesis. | (Gagliardi et al., 2020, 2017) |
| MFAP5 | Microfibril associated protein 5 | MFAP5 encode extracellular matrix proteins that play important roles in bone, blood vessels, hemostasis and the immune system. Maybe | One of the genes of a signature gene set involved in epithelial-mesenchymal transition in GBM. | (Cheng et al., 2012; Wu et al., 2019, p. 5) |

| | | | | |
|---|---|---|---|---|
| | | upregulated in invasive types of cancers. | | |
| CRIP1 | Cysteine Rich Protein 1 | Seems to have a role in zinc absorption and may function as an intracellular zinc transport protein. Gene overexpressed in prostate and pancreatic cancers. | Novel marker found to be expressed in glioma-associated brain macrophages. | (Ochocka et al., 2019; Wang et al., 2007) |
| LUM | Lumican | Lumican regulates collagen fibril organization, corneal transparency, epithelial cell migration and tissue repair. | Shown to be upregulated in GSCs and may be responsible for chemoresistance of the tumor. | (Chakravarti, 2002; Farace et al., 2015) |
| MYL9 | Myosin Light Chain 9 | The encoded protein binds calcium and is activated by myosin light chain kinase and regulates muscle contractions. It's role has been identified in colorectal and non-small cell lung carcinomas. | It is a novel biomarker associated with poor prognosis in GBM and a good determinant of its aggressiveness as it's highly expressed in recurrent GBM. | (Kruthika et al., 2019, p. 9; Luo et al., 2014, p. 9; Tan and Chen, 2014, p. 9) |

| | | | | |
|---|---|---|---|---|
| RAB34 | RAB34, Member RAS Oncogene Family | This gene encodes a protein belonging to the RAB family of proteins, which are small GTPases involved in protein transport. It is involved in hedgehog signaling and also plays a role in adhesion, invasion and migration of breast tumors. | It's overexpression is an indicator of poor prognosis and a lowered overall survival of high-grade GBM patients. | (L. Sun et al., 2018, p. 34; Wang et al., 2015, p. 34; Xu et al., 2018, p. 34) |
| TPM2 | Tropomyosin 2 | This gene encodes beta-tropomyosin, and mainly expressed in slow, type 1 muscle fibers. This gene may also be responsible for the transformation of breast epithelial cells. | The downregulation of this gene is responsible for the infiltration of the soft brain environment with the GBM tumor. | (Dube et al., 2016; Mitchell et al., 2019) |
| FEZF1 | FEZ Family Zinc Finger 1 | The encoded protein is thought to play a role in the embryonic migration of gonadotropin-releasing hormone secreting- and monoaminergic- neurons into the basal forebrain. | FEZF1-AS1 is a potential prognostic biomarker for GBM indicating poor prognosis of patients. It is a driver of GBM proliferation and infiltration, and thus, is a key oncogene. It also overcomes apoptosis by inhibiting the PI3K/AKT pathways in GSCs. | (Shimizu and Hibi, 2009; M. Yu et al., 2017; Yu et al., 2018) |

| | | | | |
|---|---|---|---|---|
| FBN2 | Fibrillin 2 | The protein encoded by this gene is a component of connective tissue microfibrils and may be involved in elastic fiber assembly. It is exclusively expressed in the peripheral nerves. It may a play role in tumorigenesis via regulation of the TGF-β pathway. | No known function in gliomagenesis, thus, further studies are needed. | (Charbonneau et al., 2003; van Loon et al., 2020) |
| SFTA1P | surfactant associated 1, lncRNA | SFTA1P is regarded as a tumor suppressor in non-small cell lung cancer. May be potential targets for therapy. | No known role in GBM, thus, studies are required. | (Huang et al., 2017; Zhang et al., 2017) |
| PDLIM4 | PDZ And LIM Domain 4 | PDLIM4, a LIM domain gene also known as RIL, is suspected to have tumor suppressor functions in myeloid diseases. Hypermethylation of this gene is a potential biomarker in prostate and breast cancers. | PDLIM4 expression is associated with imparting GBM radioresistance, and is correlated with the aggressiveness and prognosis of brain cancer. | (Feng et al., 2010; Ming et al., 2017; Tayrac et al., 2011; Vanaja et al., 2006) |

| | | | | |
|---|---|---|---|---|
| TUSC3 | Tumor Suppressor Candidate 3 | It is involved in cellular magnesium uptake, protein glycosylation and embryonic development. This gene is a candidate novel tumor suppressor gene. | Downregulation of this gene is correlated with higher grade of glioma, as it is involved in enhancing the proliferation and invasiveness of the tumor. | (X. Yu et al., 2017, p. 3; Yuan et al., 2018) |
| LOXL1 | Lysyl Oxidase Like 1 | It may be involved in developmental regulation, senescence, tumor suppression, cell growth control, and chemotaxis. | Overexpression is indicative of higher glade glioma. It is a great biomarker that can be exploited in liquid biopsy to monitor tumor progression. It is also responsible for preventing apoptosis in gliomas. | (Kim et al., 2014; M. Li et al., 2019; Yu et al., 2020) |
| SYT1 | Synaptotagmin 1 | The synaptotagmins are Ca2+ sensors responsible for vesicular trafficking and exocytosis, and triggering neurotransmitter release at the synapse. | It is a marker of the Neural subtype of GBM. More studies are required to identify its role in gliomagenesis. | (Bacaj et al., 2013; Zhang et al., 2020) |

| | | | | |
|---|---|---|---|---|
| CAV2 | Caveolin 2 | The protein is involved in essential cellular functions, including signal transduction, lipid metabolism, cellular growth control and apoptosis. This protein may function as a tumor suppressor. | CAV2 is regulated by miR-144 and enhances glioma migration and invasion via EMT. | (Liu et al., 2020, p. 2; Sowa, 2011) |
| ARHGAP 29 | Rho GTPase Activating Protein 29 | Rap1 is a small GTPase that, through effectors, regulates Rho GTPase signaling. It is a known promoter of metastasis in several types of cancers such as breast and prostate. | Knockdown of this gene is responsible for the decrease in migration of GBM. | (Bruning - Richardson et al., 2018; Kolb et al., 2020) |
| OMD | Osteomodulin | It is a marker in the human dental pulp stem cells. It may be involved in the regulation of osteogenesis. | No known association with gliomagenesis and thus, needs further investigation. | (Lin et al., 2019; Ninomiya et al., 2007) |
| XLOC_01 2003 | Novel gene | Not known. | Not known. | |
| SLC2A12 | Solute Carrier Family 2 Member 12 | SLC2A12 is involved in regulating blood urate levels. It may be a good biomarker for gastric | Not known thus further studies are needed. | (Toyoda et al., 2020; Zheng et |

| | | | | |
|---|---|---|---|---|
| | | cancer as its levels are lowered post-operation. | | al., 2020, p. 2) |
| XLOC_047367 | p53-regulated lncRNAs | Not known. | Not known. | (Jain et al., 2016) |
